# Fibrotic scar after experimental autoimmune encephalomyelitis inhibits oligodendrocyte differentiation

**DOI:** 10.1101/819128

**Authors:** Stephanie L. Yahn, Jiajun Li, Irene Goo, Han Gao, Roberta Brambilla, Jae K. Lee

## Abstract

Remyelination failure is a crucial component of disease progression in the autoimmune demyelinating disease Multiple Sclerosis (MS). The regenerative capacity of oligodendrocyte progenitor cells (OPCs) to replace myelinating oligodendrocytes is likely influenced by many aspects of the lesion environment including inflammatory signaling and extracellular matrix (ECM) deposition. These features of MS lesions are typically attributed to infiltrating leukocytes and reactive astrocytes. Here we demonstrate that fibroblasts also contribute to the inhibitory environment in the animal model of MS, experimental autoimmune encephalomyelitis (EAE). Using Col1α1^GFP^ transgenic mice, we show that perivascular fibroblasts are activated in the spinal cord at EAE onset, and infiltrate the parenchyma by the peak of behavioral deficits where they are closely associated with areas of demyelination, myeloid cell accumulation, and ECM deposition. We further show that both fibroblast conditioned media and fibroblast ECM inhibit the differentiation of OPCs into mature oligodendrocytes. Taken together, our results indicate that the fibrotic scar is a major component of EAE pathology that leads to an inhibitory environment for remyelination, thus raising the possibility that anti-fibrotic mechanisms may serve as novel therapeutic targets for MS.

## Introduction

In multiple sclerosis (MS), proper remyelination and functional recovery is chronically impaired. The failure of remyelination despite the presence of oligodendrocyte progenitor cells (OPCs; NG2 glia) within and around demyelinated areas is suggestive of an inhibitory environment. The demyelinating insult results in a multi-cellular response that includes inflammatory signaling and excessive extracellular matrix (ECM) deposition, which may inhibit the regenerative capacity of OPCs. To therapeutically target the production of inhibitory cues and create a more permissive environment for remyelination, the source of such cues must be elucidated. The current FDA approved therapeutic interventions for MS solely target immune cell activation and inflammatory signaling, which slow the progression of the disease. There is a clear need to develop therapies capable of modulating the responses of other cell types in the lesion as well as the excessive accumulation of ECM in order to promote recovery. Establishing the cellular origin of deposited ECM is difficult, however, since ECM antibodies label the extracellular environment rather than the cells themselves. Given that astrocytes are regarded as the major producers of ECM in the CNS, they have received the most attention for their role in scar formation (Hibbits, Yoshino, Le, & Armstrong, 2012; Lau, Cua, Keough, Haylock-Jacobs, & Yong, 2013; Stoffels et al., 2013; Stoffels, Hoekstra, Franklin, Baron, & Zhao, 2015). However, other cellular contributions to ECM deposition and the regulation of the extracellular environment are not fully understood.

There is increasing evidence that fibrotic scarring contributes to the pathology of several types of CNS injury (Dias & Goritz, 2018; Fernandez-Klett & Priller, 2014). We previously demonstrated that perivascular fibroblasts are the source of the fibrotic scar after spinal cord injury (SCI) (Soderblom et al., 2013) and that fibrotic scar formation is mediated by hematogenous macrophages that migrate to the injury site (Zhu et al., 2015). The neuroinflammatory mechanism of fibrosis after SCI raised the possibility that fibrosis may be a pathology common to many neuroinflammatory diseases including multiple sclerosis. Much of what is known about fibrosis comes from studies of peripheral tissues, which have shown that fibrosis occurs when normal wound healing mechanisms are not properly regulated resulting in a state of uncontrolled tissue remodeling that can prevent proper functional regeneration. Fibroblasts play a pivotal role in the alteration of the injury environment through the production of ECM (such as collagens, fibronectin, laminin, proteoglycans, and glycoproteins), cytokines, and growth factors. Interestingly, fibrotic ECM accumulation, such as collagen, has been documented in human MS lesions (Mohan et al., 2010; Sobel, 1998; Sobel & Ahmed, 2001; van Horssen, Bo, Dijkstra, & de Vries, 2006) as well as animal models of MS (Haist, Ulrich, Kalkuhl, Deschl, & Baumgartner, 2012; Teesalu, Hinkkanen, & Vaheri, 2001), but the cellular origins and functional significance of fibrosis in MS has not been clearly established.

The goal of this study was to investigate the temporospatial profile of fibrotic scar formation and its contribution to oligodendrocyte differentiation in a mouse model of MS, experimental autoimmune encephalomyelitis (EAE). Using transgenic mice in which fibroblasts are labeled with green fluorescent protein (GFP), we demonstrate that fibroblasts are a major cellular component of EAE lesions and are closely associated with fibrous ECM deposition in demyelinated areas. To test whether fibroblasts influence OPC differentiation, we cultured OPCs with fibroblasts and show that fibroblasts do not promote oligodendrogenesis. Furthermore, we demonstrate that OPC differentiation is inhibited by fibroblast conditioned media and fibroblast ECM. Overall, we present the first systematic study of the fibrotic scar in EAE lesions and provide evidence suggesting that fibroblasts contribute to an inhibitory environment for OPC differentiation. Previous studies have alluded to the presence of fibrosis in MS post-mortem tissue which underscores the importance of further elucidating the role of the fibrotic scar in MS lesion pathology. We have demonstrated the cellular source of fibrosis in a commonly used animal model of MS, which permits more mechanistic studies of fibrosis after EAE in the future.

## Materials and methods

### EAE induction

Col1α1^GFP^ mice in C56BL/6 background were kindly donated by Dr. David Brenner, University of California San Diego, La Jolla, CA (Yata et al., 2003). Male Col1α1^GFP^ mice (2-4 months old) received an injection of pertussis toxin (i.p., 500ng/mouse on days 0 and 2; List Biological Laboratories, Cat.# 181) and an injection of MOG_35-55_ peptide (s.c., 300ng/mouse on day 1; BioSynthesis Cat# 12668-01) emulsified in Complete Freund Adjuvant (7.5mg/mL M. Tuberculosis H37RA (Fisher Cat# DF3114338) in adjuvant (Fisher Cat# DF0639606)). Motor deficits were recorded daily starting on day 7 until the end of the experiment. The scoring criteria was based on a 6 point scale as follows: 0-no apparent symptoms, 1-loss of tail tone, 2-flaccid tail, 3-complete hindlimb paralysis, 4-complete forelimb paralysis, 5-moribund, 6-dead. Half-points were also used to better represent the transition to paralysis (i.e. 2.5-hindlimb weakness). Mice were sacrificed at disease onset (2 consecutive days scoring at least a 2), acute disease (7-9 days after onset) and chronic disease (60 days post induction). Mice that did not reach a score of at least 2.5 were excluded from analyses. All animal procedures were approved by the University of Miami Institutional Animal Care and Use Committee.

### Histology

At onset, acute and chronic time points after EAE induction (n=5 per group), mice were perfused transcardially with 4% paraformaldehyde (PFA). Spinal cord and optic nerves were dissected, postfixed for 24 hours in 4% PFA, cryoprotected in 30% sucrose overnight and then embedded in OCT compound. Serial transverse cryosections (16um) of the thoracic spinal cord were thaw mounted onto Superfrost Plus slides, washed with PBS-0.3% TritonX-100 (PBS-T), blocked in 5% goat serum (in PBS-T) for 1hr at room temperature and then incubated overnight at 4°C with antibodies in blocking solution against GFP (Abcam Cat# ab13970, RRID:AB_300798, 1:2000), CD11b (Thermo Fisher Scientific Cat# RM2800, RRID:AB_10375863, 1:500), fibronectin (Millipore Cat# AB2033, RRID:AB_2105702, 1:500), GFAP (Dako Cat# Z0334, RRID:AB_10013382, 1:1000), NG2 (Millipore Cat# AB5320, RRID:AB_91789, 1:200), or PDGFRβ (Abcam Cat# ab32570, RRID:AB_777165, 1:500). After primary antibody incubation, sections were washed in PBS-T and then incubated in the appropriate Alexa Fluor secondary antibodies (Invitrogen, 1:500) for 1hr at room temperature. After washing off the secondary antibody with PBS-T, sections were stained with FluoroMyelin (Thermo Fisher Scientific Cat# F34652, RRID:AB_2572213, 1:300) or DAPI and coverslipped in Fluoromount G mounting solution. Images were collected with a Nikon Eclipse Ti epifluorescent microscope.

### Cell cultures

Fibroblasts were cultured from cortical meninges of postnatal day 5 CD1 mouse pups. The brain was dissected from the skull and meninges were carefully removed from the cortex using fine-tipped forceps and collected into cold HBSS (Thermo Fisher Scientific Cat# 14175-145). The meninges were digested in trypsin (VWR Cat# 45000-664) with collagenase (Worthington Biochemical Cat# LS0004174) for 30 minutes at 37°C with intermittent trituration. The cells were rinsed with DMEM (Sigma-Aldrich Cat# 11995-073) supplemented with 10% FBS (Thermo Fisher Scientific Cat# 26140-079) and filtered through a 70µm cell strainer (VWR Cat# 21008-952) before culturing in a T-75 tissue culture flask (VWR Cat# 82050-856) coated with poly-D-lysine (PDL, Millipore Cat# A-003-E). Fibroblasts were split once before generating co-cultures, ECM coated wells, or conditioned media.

Astrocytes were purified from primary mixed glial cultures using an anti-GLAST MicroBead Kit (Miltenyi Biotec Cat# 130-095-826). Mixed glial cultures were generated from post-natal day 5 CD1 mouse pup cortices. After the removal of the meninges, brains were collected into ice cold HBSS and minced using a razor blade. The brains were digested in trypsin and enzyme A from the Neural Tissue Dissociation Kit (Miltenyi Biotec Cat# 130-092-628) for 30 minutes at 37°C with intermittent trituration. The cells were rinsed with DMEM supplemented with 10% FBS. The mixed cell suspension was cultured in a T-75 tissue culture flask coated with PDL. When an astrocyte monolayer formed in the flask after approximately seven days, the cells were trypsinized from the flask and rinsed with DMEM with 10% FBS. The cells were then rinsed with DPBS (Thermo Fisher Scientific Cat# 14287-080) with 0.5% BSA (Sigma-Aldrich Cat# A9576) prior to purification of astrocytes following the manufacturer’s instructions from the anti-GLAST Microbead Kit (Miltenyi Biotec Cat# 130-095-826). The purified astrocytes were cultured in a T-75 tissue culture flask coated with PDL and maintained in DMEM with 10% FBS.

OPCs were isolated and purified from postnatal day 5 CD1 mouse cortices. Brains were collected into ice cold HBSS and minced using a razor blade. The brains were digested using the Neural Tissue Dissociation Kit-P from Miltenyi Biotec following the manufacturer’s protocol. OPCs were purified from the cortical cell suspension using a PDGFR-α Microbead Kit (Miltenyi Biotec Cat# 130-101-502). OPCs were collected into OPC media (DMEM-F12 (Thermo Fisher Scientific Cat# 11320-033), supplemented with 1% B27 (Thermo Fisher Scientific Cat# 17504-044), 1% N2 (Thermo Fisher Scientific Cat# 17502-048), 1% penicillin/streptomycin (Thermo Fisher Scientific Cat# 15140122), 40 ng/mL bFGF (Sigma-Aldrich Cat# F0291), and 20 ng/µL PDGF-AA (Thermo Fisher Scientific Cat# PHG0035)).

### OPC co-cultures with fibroblasts and/or astrocytes

To generate cell monolayers, fibroblasts or astrocytes were split into PDL-coated 48-well plates (VWR Cat# 82051-004) five days prior to the addition of OPCs to allow for the formation of a confluent cell monolayer. For the fibroblasts with astrocytes condition, astrocytes and fibroblasts were mixed together in equal parts prior to plating. To label and track OPCs, purified primary OPCs were pre-labeled with cytopainter-green dye (Abcam Cat# ab176735) for 15 minutes prior to plating them onto cell monolayers or PDL. 3×10^4^ cytopainter labeled OPCs were plated in each condition and maintained in OPC media. To test the effect of growth factor withdrawal, media was changed after one day to complete media lacking PDGF-AA and bFGF. To stimulate OPC differentiation, media was changed after two days to complete media lacking PDGF-AA and bFGF supplemented with 40 ng/mL T_3_ (Sigma-Aldrich Cat# T5516), 1 ng/mL CNTF (Sigma-Aldrich Cat# C3835), and 50 µg/mL insulin (Sigma-Aldrich Cat# I2643).

To test the proliferation of OPCs on the second day in vitro, approximately 16 hours after the initiation of growth factor withdrawal, 10 µM EdU was added to the media and cells were fixed 8 hours later using cold 4% PFA. EdU was visualized using a Click-iT EdU Alexa Fluor 647 Imaging Kit (Thermo Fisher Scientific Cat# C10639). To test the amount of cell death on the second day in vitro approximately 24 hours after the initiation of growth factor withdrawal, a LIVE/DEAD fixable cell stain (Thermo Fisher Scientific Cat# L34973) was added to the media for 30 minutes prior to fixation with cold 4% PFA.

### OPC cultures with conditioned media

Conditioned media from fibroblasts, astrocytes, or OPCs was generated by plating 3×10^4^ cells per cm^2^ in PDL-coated 6-well plates. The cells were allowed to adhere, and after one hour they were rinsed three times with HBSS followed by the addition of 2mL per well of DMEM-F12 supplemented with 1% N2, 1% B27, 0.01% BSA, and penicillin-streptomycin. The media was collected 24 hours later and filtered through a 0.2 micron syringe filter to remove cellular debris and then frozen at −80°C until use. OPCs were plated at 3×10^4^ cells per cm^2^ in PDL-coated 48-well plates in OPC media (mentioned above). After 24 hours, the OPC media was changed to conditioned media supplemented with 40 ng/mL T_3_ (Sigma-Aldrich Cat# T5516), 1 ng/mL CNTF (Sigma-Aldrich Cat# C3835), and 50 µg/mL insulin (Sigma-Aldrich Cat# I2643). The media was refreshed two days later, and on the next day the plates were fixed with 4% PFA.

### OPCs cultured on fibroblast ECM

Fibroblast ECM-coated wells were generated by culturing fibroblasts in 48-well plates coated with PDL for five days prior decellularization using a non-enzymatic cellular dissociation buffer (Thermo Fisher Scientific Cat# 13150-016). Decellularization was confirmed by bright-field microscopy and later by DAPI staining. ECM wells were washed with HBSS and used immediately for OPC cultures. OPCs were plated at 3×10^4^ cells per cm^2^ in OPC media. After 24 hours, the OPC media was changed to differentiation media (DMEM-F12 supplemented with 1% N2, 1% B27, 0.01% BSA, 40 ng/mL T_3_, 1 ng/mL CNTF, and 50 µg/mL insulin, and penicillin-streptomycin). The media was refreshed two days later, and the plates were fixed with 4% PFA after four days of differentiation.

### Immunocytochemistry

Cell cultures were fixed with 4% PFA for 15 minutes followed by three washes with PBS. The cells were washed with PBS-T, blocked with 5% goat serum (in PBS-T) for 1hr at room temperature and then incubated overnight at 4°C with antibodies in blocking solution against MBP (Millipore Cat# MAB386 RRID:AB_94975, 1:500), fibronectin (Millipore Cat# AB2033, RRID:AB_2105702, 1:500), GFAP (Dako Cat# Z0334, RRID:AB_10013382, 1:1000), or NG2 (Millipore Cat# AB5320, RRID:AB_91789, 1:200). After primary antibody incubation, cells were washed in PBS-T and then incubated in the appropriate Alexa Fluor secondary antibodies (Invitrogen, 1:500) for 1hr at room temperature. After washing off the secondary antibody with PBS-T, cells were stained with DAPI.

### Quantification

For Figures 1–3, area contours of GFP positive (infiltrating fibroblasts), CD11b positive (dense accumulation of round myeloid cells), fibronectin positive, and fluoromyelin negative regions were measured using the polygon tool in ImageJ. Four full transverse sections of thoracic spinal cord were quantified per animal for each time point. The contoured areas were normalized to total white matter area (area contours of white matter) per section. Fibroblast density was quantified by counting the number of GFP positive cells with DAPI positive nuclei within the parenchyma (excluding the meninges and obvious blood vessels) and normalized to white matter area.

**Figure 1.**
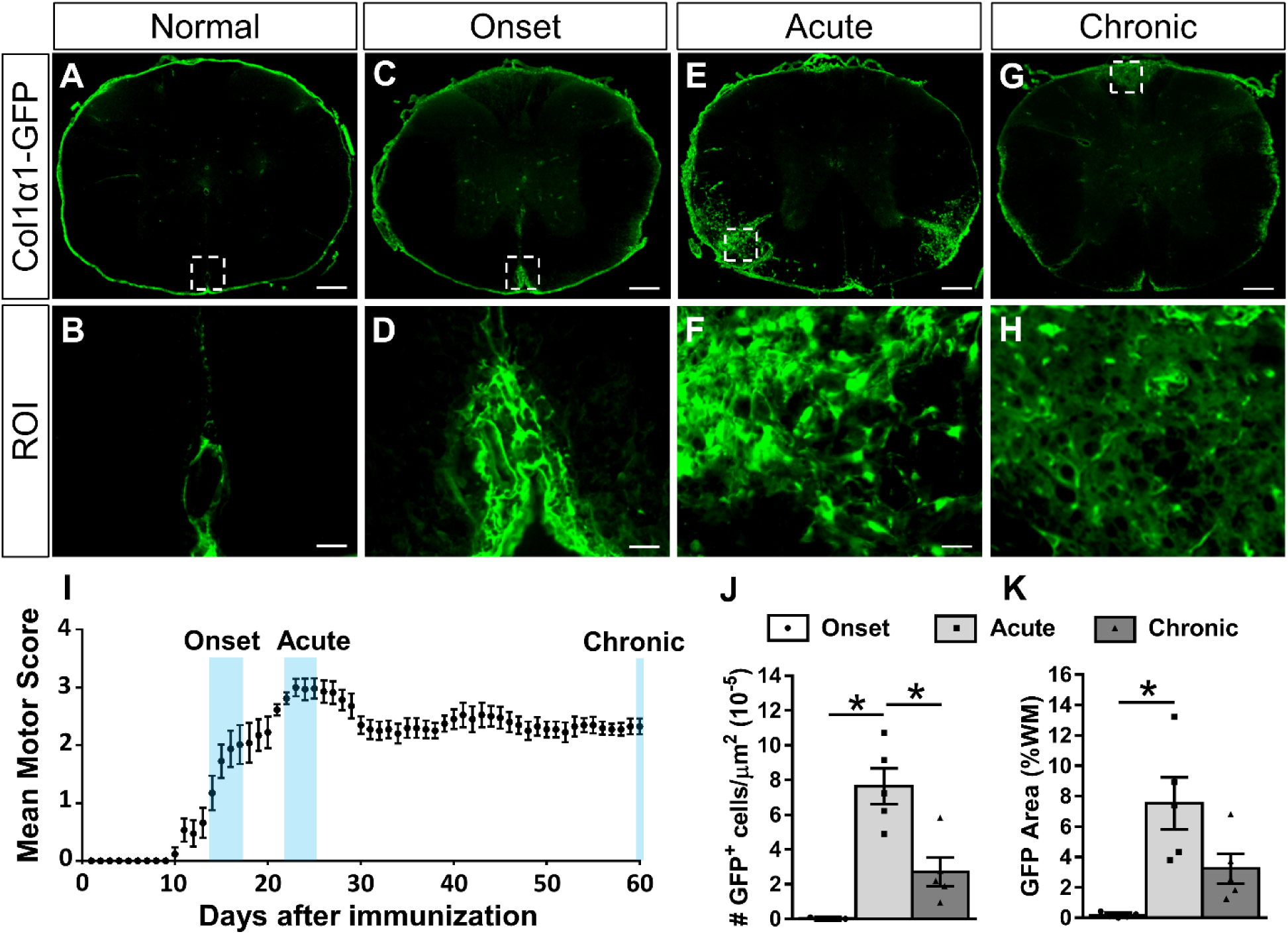
Fibroblasts infiltrate the white matter acutely after EAE and then partially resolve over time. Cross section of thoracic spinal cord from a normal Col1α1^GFP^ mouse shows fibroblasts in the meninges and around large diameter blood vessels (A-B). At the onset of motor deficits after EAE induction (I), fibroblasts are enlarged but primarily remain perivascular/meningeal (C-D). During acute disease when motor deficits have reached a maximum (I), fibroblasts display peak infiltration into the white matter parenchyma (E-F, J-K). At 60 days post induction when motor scores have plateaued chronically (I), fibroblasts in the white matter are less reactive (G-H) and fewer in number (J). Scale bars = 200μm (A, C, E, G) and 25μm (ROI). n=5 biological replicates for each time point. *p<0.05 one-way ANOVA with Bonferonni corrections for multiple comparisons, error bars represent SEM.

**Figure 2.**
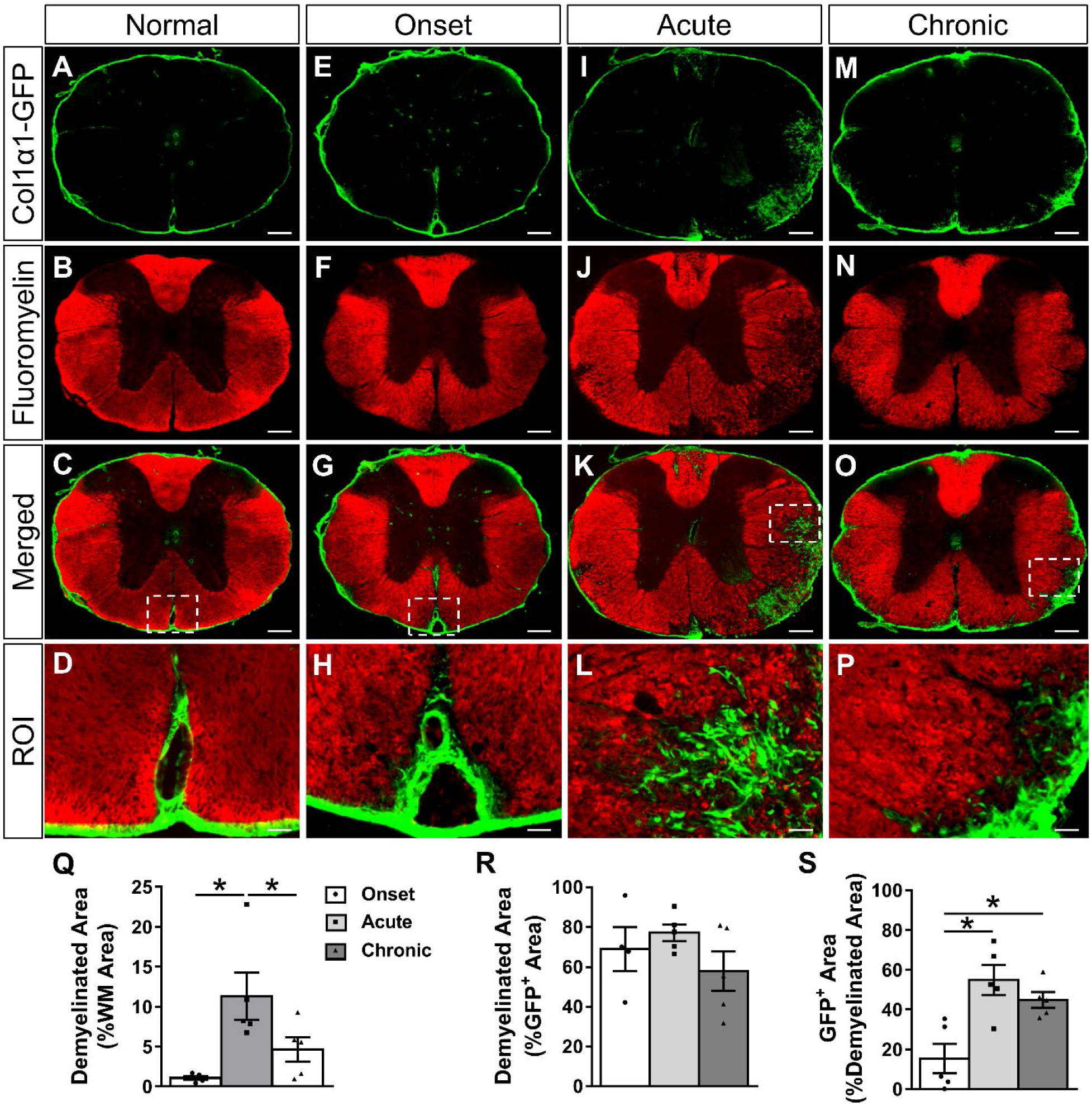
Fibrotic regions are closely associated with demyelination acutely and chronically after EAE. Cross section of thoracic spinal cord from a normal Col1α1^GFP^ mouse shows compact myelin (A-D) that begins to lose its structure and small demyelinated areas appear by the onset of EAE (E-H). By the acute disease phase, the percent of white matter that is demyelinated increases (I-L, Q), and fibroblasts are mainly localized to demyelinated areas (R). As the disease progresses from the onset to acute phase, the demyelinated area becomes increasingly filled with fibroblasts, and chronic demyelinated areas continue to contain fibroblasts (S). The GFP^+^ area includes perivascular GFP signal in demyelinated areas. Boxed areas in C, G, K, O are regions of interest (ROI) that are enlarged in D, H, L, P. Scale bars = 200μm (first three rows) and 25μm (ROI). n=5 biological replicates for each time point. *p<0.05 one-way ANOVA with Bonferonni corrections for multiple comparisons, error bars represent SEM.

**Figure 3.**
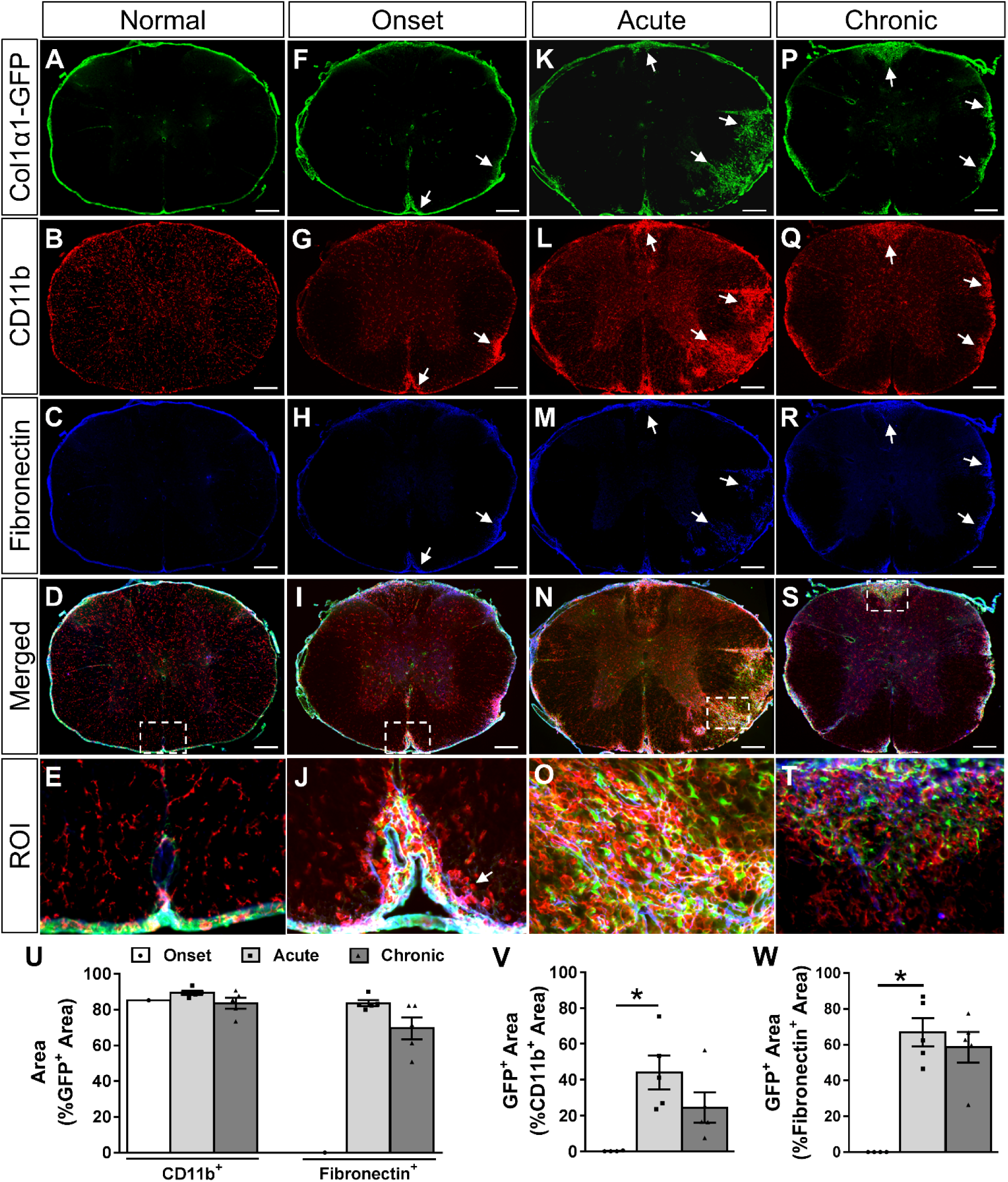
Fibrotic regions are closely associated with myeloid cell accumulation and fibronectin deposition acutely and chronically after EAE. Cross section of thoracic spinal cord from a normal Col1α1^GFP^ mouse shows fibroblasts (A) and fibronectin (C) exclusively in the meninges and around blood vessels. At disease onset, CD11b^+^ myeloid cells (G) and fibronectin (H) begin to accumulate in the white matter (arrows). During acute disease, fibroblasts infiltrate the parenchyma at sites of CD11b^+^ myeloid cell accumulation (K, L, N, O). Over 80% of the fibrotic GFP^+^ area has an accumulation of CD11b^+^ myeloid cells at all stages of disease (U). At the onset of physical symptoms, CD11b^+^ myeloid cells begin accumulating in the white matter before the infiltration of fibroblasts, but approximately 50% of the dense CD11b^+^ regions are filled with fibroblasts by the acute stage of disease (V). Similarly, about 80% of the fibroblastic GFP^+^ regions are fibronectin^+^ during acute and chronic disease (U), and about 60% of the fibronectin^+^ areas have fibroblasts (W). Fibroblasts were observed in the white matter of only one animal at onset (U). Boxed areas in D, I, N, S are regions of interest (ROI) that are enlarged in E, J, O, T. Onset n=4, acute and chronic n=5 (biological replicates). Scale bars = 200μm (first 4 rows) and 25μm (ROI). *p<0.05 one-way ANOVA with Bonferonni corrections for multiple comparisons, error bars represent SEM.

For Figures 4–6, cytopainter labeled cells with DAPI^+^ nuclei were manually counted using ImageJ software in four non-overlapping fields taken at the center of each well with a 20x objective of a Nikon Eclipse Ti epifluorescent microscope. The percent of cytopainter labeled OPCs that proliferated (EdU^+^), were dead or dying (positive for the dead cell stain) or differentiated into oligodendrocytes (MBP^+^) were manually counted using ImageJ. To quantify the percent of OPCs that differentiated on fibroblast ECM compared to PDL in Figure 7, MBP^+^ cells with DAPI^+^ nuclei were manually counted using ImageJ in four non-overlapping fields taken with a 20x objective. To quantify the percent of OPCs that differentiated in conditioned media in Figure 8, MBP^+^ cells with DAPI^+^ nuclei were manually counted using ImageJ in nine non-overlapping fields taken at 10x using an ArrayScan high-content imaging system. For each *in vitro* assay, three separate wells were used as technical replicates, and the experiment was repeated three times (i.e. ECM-coated plates for each experimental replicate were generated from a separate fibroblast isolation, and conditioned media for each experimental replicate was generated from separate cell isolations). Experimenters were blind to conditions while quantifying cells for the conditioned media and ECM experiments. The conditions were obvious for the co-culture experiments (fewer nuclei in the PDL wells; elongated and enlarged nuclei in the fibroblast wells), so experimenters were not able to maintain a blinded approach for quantifications.

**Figure 4.**
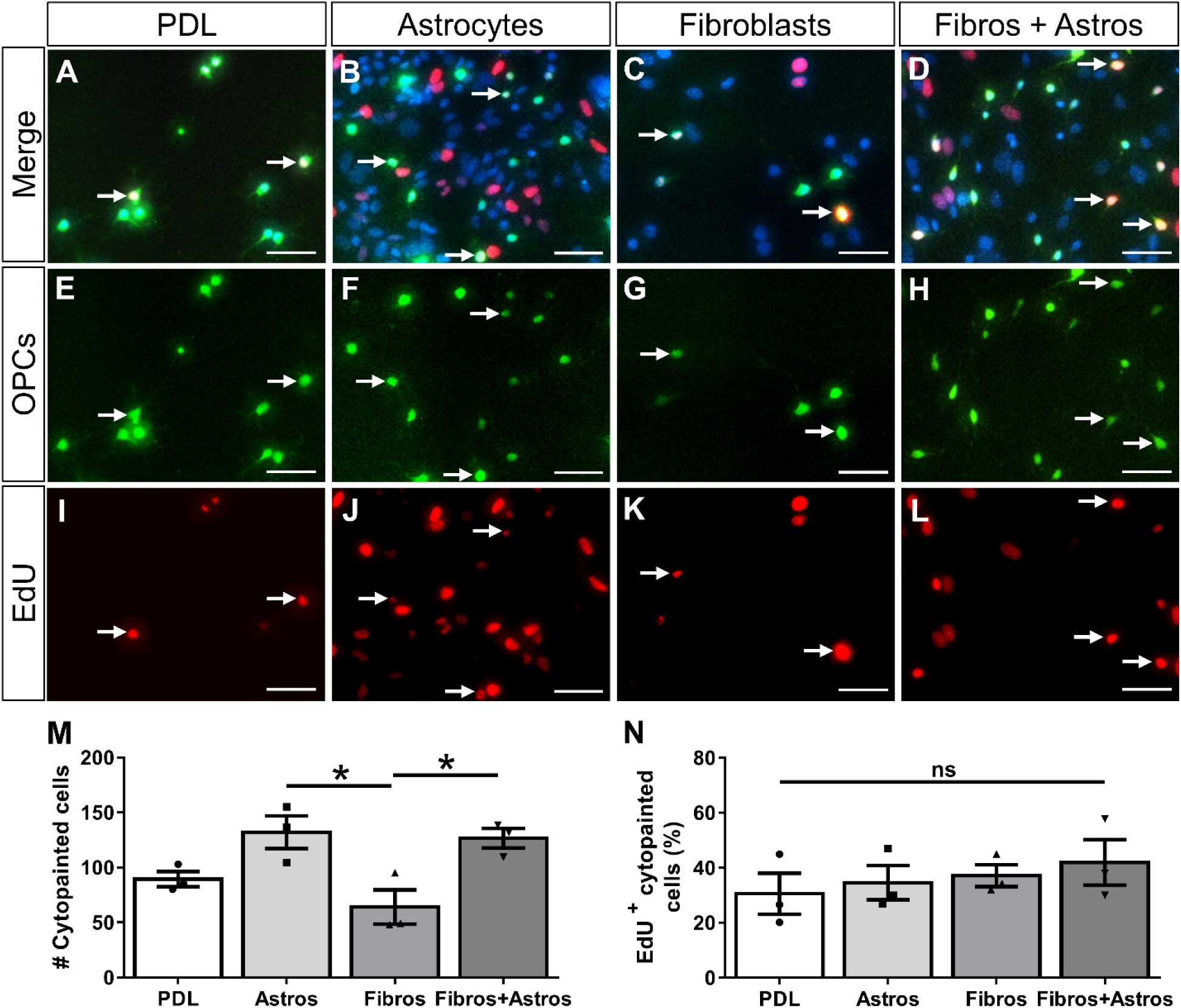
Fibroblasts do not affect the proliferation of OPCs. Primary oligodendrocyte progenitor cells (OPCs) pre-labeled with cytopainter-green dye were cultured on PDL or monolayers of astrocytes, fibroblasts, or a mix of astrocytes and fibroblasts for 2 days (A-H). OPC proliferation was assessed by EdU incorporation (I-L, arrows indicate example OPCs that have incorporated EdU). Fewer cytopainted OPCs were observed on fibroblast monolayers compared to astrocyte monolayers or mixed astrocyte+fibroblast monolayers (M). The percent of proliferating cytopainted OPCs was not different between substrates (N). n=3 biological replicates per group, *p<0.05 one-way ANOVA with Bonferonni corrections for multiple comparisons, error bars represent SEM. Scale bars = 50μm. ns=not significant

**Figure 5.**
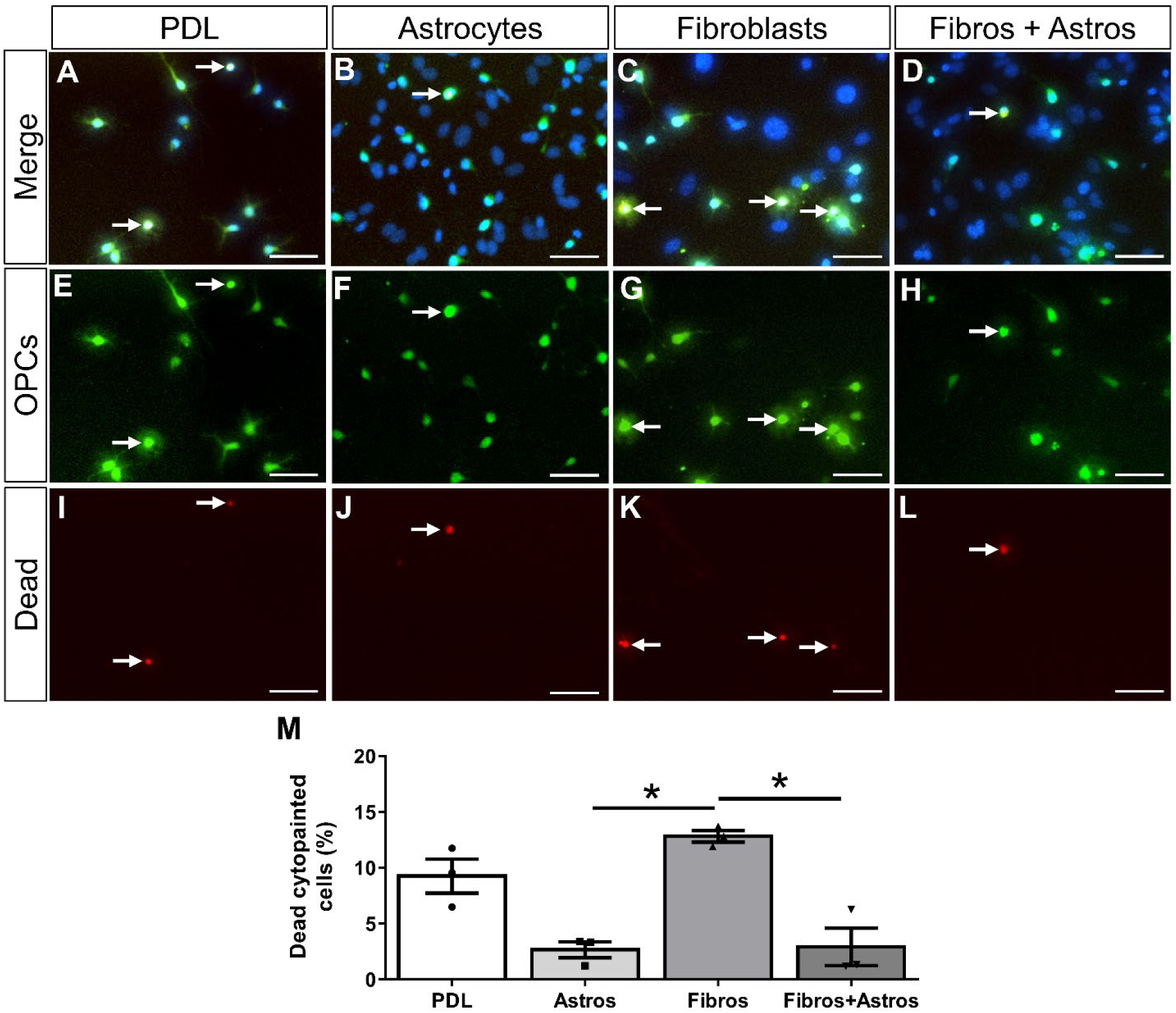
Fibroblasts do not support OPC survival. Primary OPCs pre-labeled with cytopainter-green dye were cultured on PDL or monolayers of astrocytes, fibroblasts, or a mix of astrocytes and fibroblasts for 2 days (A-H). OPC death was assessed by the incorporation of a cell-impermeable dye into cells with broken membranes (I-L, arrows indicate OPCs that have incorporated dead cell stain). The percent of cytopainted OPCs that were dead or dying was significantly more on PDL and fibroblast monolayers compared to astrocyte monolayers or mixed astrocyte+fibroblast monolayers (M). n=3 biological replicates per group, *p<0.05 One-way ANOVA with Bonferonni corrections for multiple comparisons, error bars represent SEM. Scale bars = 50μm.

**Figure 6.**
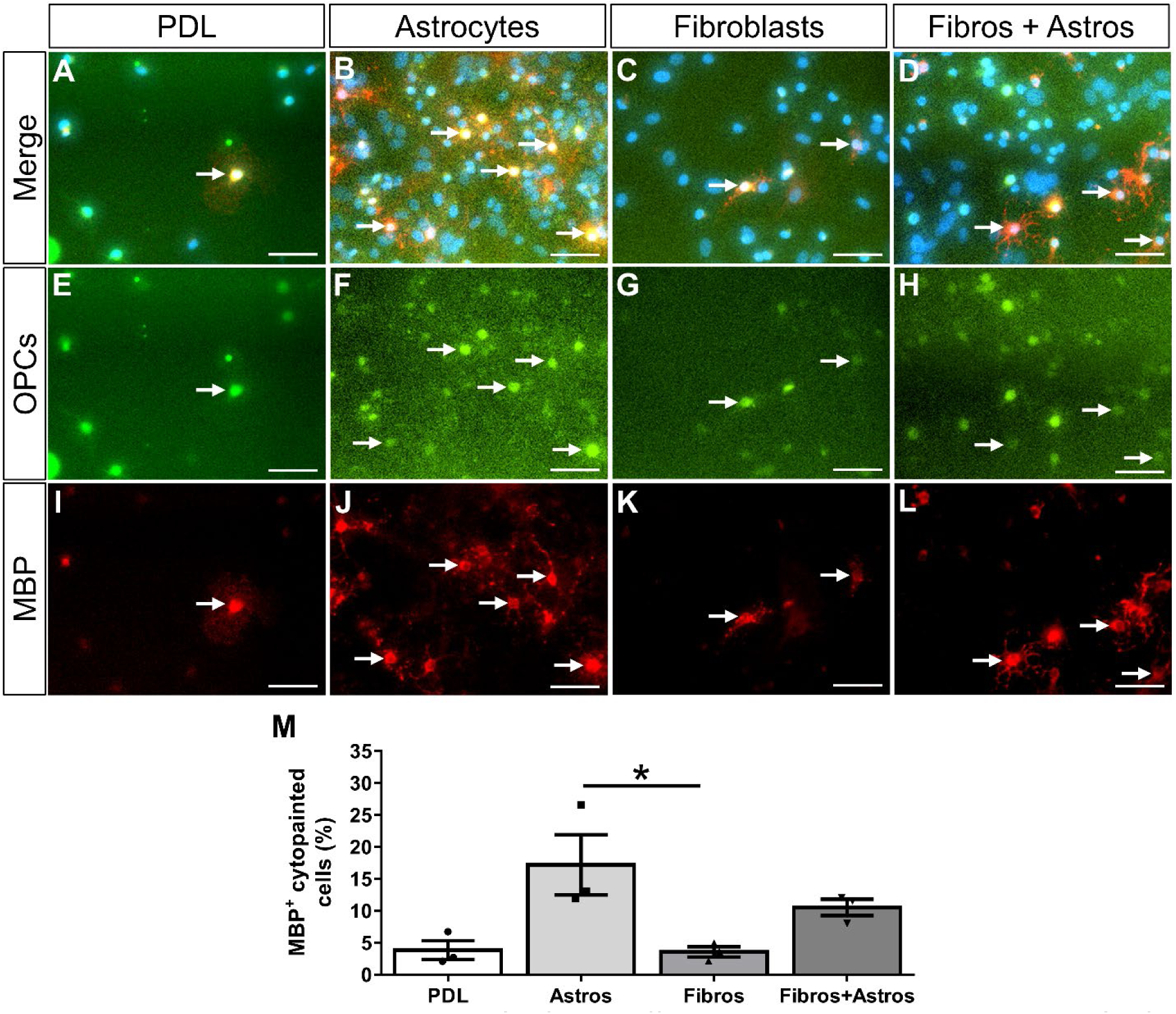
Fibroblasts do not induce OPCs to differentiate into oligodendrocytes. OPCs pre-labeled with cytopainter-green dye were cultured on PDL or monolayers of astrocytes, fibroblasts, or a mix of astrocytes and fibroblasts for 4 days (A-H). OPC differentiation was assessed by the expression of MBP (I-L, arrows indicate example OPCs that have differentiated into MBP+ oligodendrocytes). Similar numbers of cytopainted OPCs were observed across all substrates (data not shown). The percent of cytopainted OPCs that differentiated into oligodendrocytes after 3 days of PDGFA and bFGF removal was significantly more on astrocyte monolayers compared to fibroblast monolayers or PDL (M). n=3 biological replicates per group, *p<0.05 One-way ANOVA with Bonferonni corrections for multiple comparisons, error bars represent SEM. Scale bars = 50μm.

**Figure 7.**
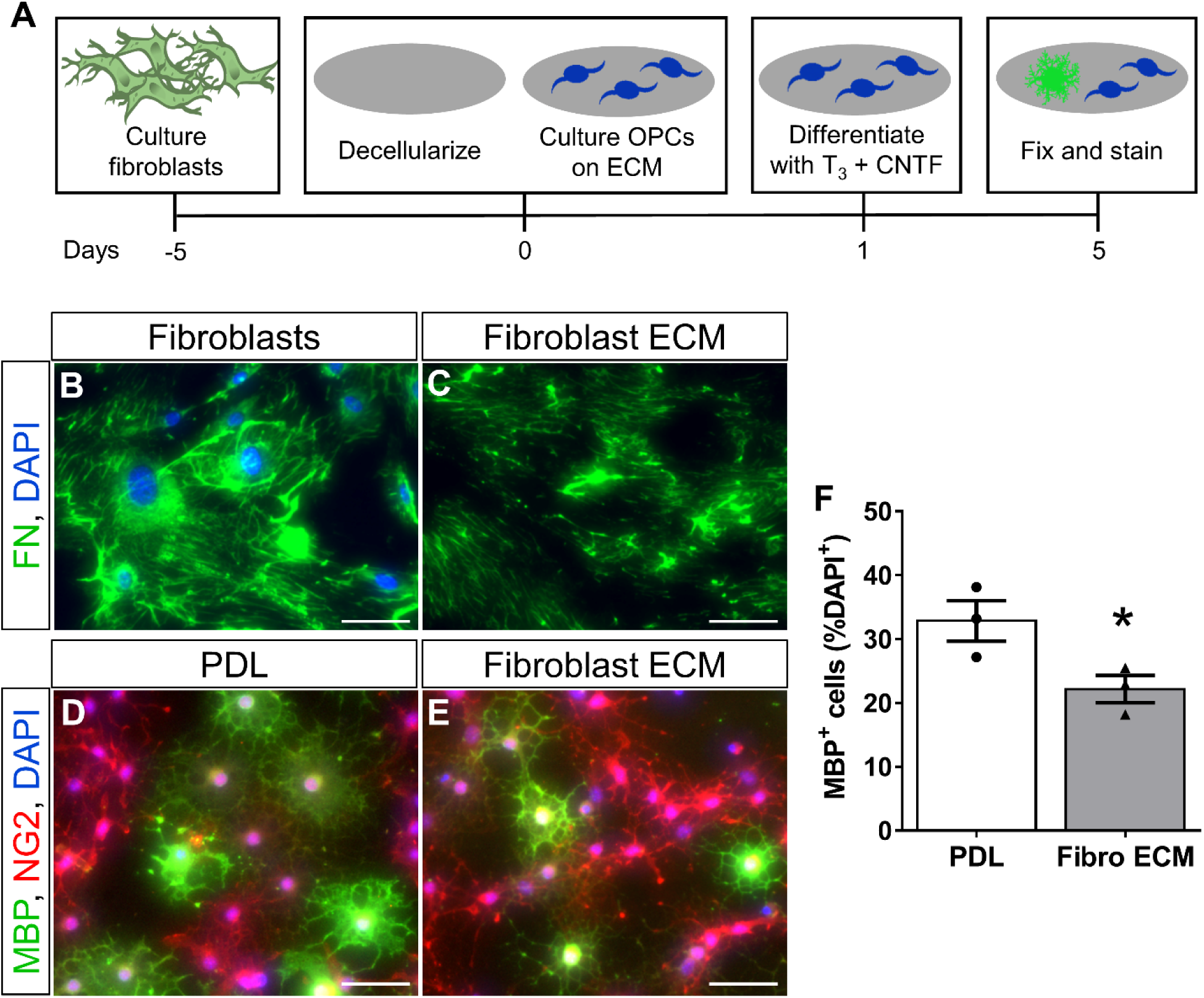
Fibroblast ECM inhibits oligodendrogenesis in vitro. OPCs are cultured on PDL or PDL with secreted fibroblast ECM, and then stimulated to differentiate using T_3_ and CNTF (A). Fibroblasts deposit ECM, including fibronectin (FN), which remain in culture after removing fibroblasts by non-enzymatic cellular dissociation (B-C). The stimulated differentiation of OPCs is inhibited on fibroblast ECM compared to PDL (D-F). A greater percentage of MBP^+^ oligodendrocytes was observed on PDL compared to fibroblast ECM. MBP^−^ cells were NG2^+^ progenitors. n=3 biological replicates per group, *p<0.05 Student’s t-test, error bars represent SEM. Scale bars = 50μm.

**Figure 8.**
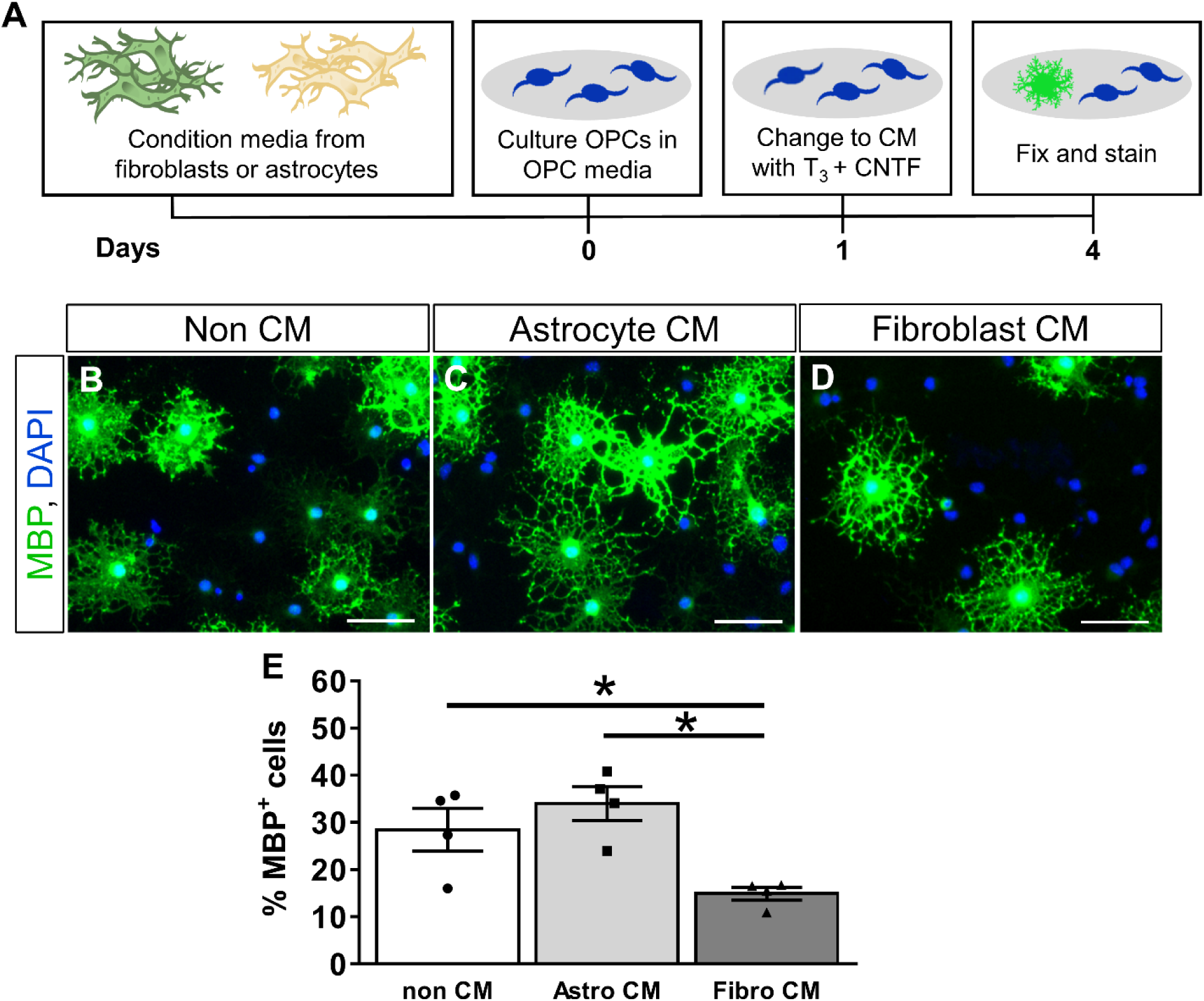
Fibroblast conditioned media inhibits oligodendrogenesis in vitro. OPCs are stimulated to differentiate in non-conditioned media or conditioned media from fibroblasts or astrocytes (A). Fewer OPCs differentiate into MBP^+^ oligodendrocytes in the presence of fibroblast conditioned media as compared to non-conditioned media or astrocyte conditioned media. (B-E). n=3 biological replicates per group, error bars represent SEM. Scale bar = 50μm.

## Results

### EAE leads to a fibrotic scar in the white matter that partially resolves over time

In the normal spinal cord (Fig. 1A, B), Col1α1^GFP^ fibroblasts reside in the meninges surrounding the spinal cord as well as in the perivascular space around large diameter blood vessels (Virchow-Robin space) as they enter or exit the spinal cord as we have previously described (Soderblom et al., 2013). To determine the response of perivascular/meningeal fibroblasts after EAE, we induced EAE in Col1α1^GFP^ transgenic mice and examined the thoracic spinal cord at onset, acute, and chronic time points (Fig. 1). At 14-17 days after EAE induction when motor deficits first begin to appear (i.e. onset, Fig. 1I), Col1α1^GFP^ fibroblasts appeared larger and reactive with more intense GFP fluorescence, but they remained largely within their perivascular and meningeal locations (Fig 1C, D). At 22-25 days after EAE induction when there were maximum motor deficits (i.e. acute disease), fibroblasts displayed peak infiltration into the white matter parenchyma and large areas of the white matter were observed as fibrotic regions (about 8% of the white matter, Fig. 1E, F, J, K). The infiltrating Col1α1^GFP^ fibroblasts were PDGFRβ positive (Supplementary Fig. 1), consistent with previous reports (Soderblom et al., 2013). At 60 days after EAE induction when the motor deficits have plateaued (i.e. chronic disease, Fig. 1G, H, I), there was a decrease in the number of fibroblasts as well as the area covered by fibroblasts (about 3%) (Fig. 1J-K), suggesting a partial resolution of the fibrotic scar.

Since MS often leads to optic neuritis, we also assessed the optic nerve and found a similar fibroblast response to what we observed in the spinal cord. The number of fibroblasts observed in the optic nerve was greatest during the acute phase, which corresponded to an elevated number of CD11b^+^ myeloid cells (Supplementary Fig. 2). In summary, our data demonstrate that Col1α1^GFP^ fibroblasts begin to enter the white matter after EAE at the onset of motor deficits, reach a peak of infiltration during the acute phase of disease (greatest functional deficit), and partially resolve over time as the disease progresses chronically.

### Fibrotic scar is demyelinated after EAE

To investigate the relationship between the fibrotic scar and areas of demyelination, we used Fluoromyelin to detect myelin in spinal cord tissue sections of Col1α1^GFP^ mice after EAE. In normal spinal cord (Fig. 2A-D), fibroblasts were restricted to the vasculature and were not present in myelinated areas of the spinal cord. In general, the temporal pattern of demyelination (Fig. 2Q) corresponded very closely to the pattern of fibrosis. During the onset phase of EAE, small areas of demyelination were observed, but these regions were largely devoid of fibroblasts (Fig. 2E-H, S). By the acute and chronic phases, about 50% of the demyelinated areas were filled with fibroblasts (Fig. 2I-P, S). Importantly, most fibrotic areas were demyelinated (about 80%) at the acute phase (Fig. 2I-L, R), suggesting a strong association between fibrosis and demyelination. Previous studies have implicated astrogliosis in the chronic demyelinated pathology of MS and EAE (Mayo et al., 2014; Stoffels et al., 2013). To compare astrogliosis and fibrosis in demyelinated areas, we immunostained Col1α1^GFP^ tissue sections for GFAP. Areas of astrogliosis (strong GFAP immunoreactivity) were closely associated with Col1α1^GFP^ fibrotic regions (Supplementary Fig. 4), suggesting that fibroblasts may also be associated with pathological mechanisms that have been previously attributed to astrocytes in EAE lesions. Taken together, our data demonstrate a close association between fibrosis and demyelination pathologies.

### Myeloid cell accumulation and extracellular matrix deposition in the fibrotic scar

To investigate the relationship between myeloid cell accumulation and fibroblast infiltration after EAE, we immunostained thoracic spinal cord tissue sections from Col1α1^GFP^ mice for myeloid cells using an antibody against CD11b. At the onset of EAE, inflammatory regions dense with round CD11b^+^ cells in the white matter were generally devoid of fibroblasts, suggesting that the inflammatory response precedes fibroblast infiltration into the CNS parenchyma (Fig. 3F, G, J). During the acute phase, a further increase in myeloid cell accumulation corresponded to fibroblast infiltration into the white matter (Fig. 3K, L, O). At this time, about 45% of CD11b^+^ inflammatory regions were covered with fibroblasts which decreased to about 25% in the chronic disease state (Fig. 3V). Conversely, approximately 85-90% of the GFP^+^ fibroblastic regions were associated with inflammatory regions during all phases of EAE (Fig. 3U), suggesting a close association of the fibrotic response to inflammation.

Previous studies have reported dense fibronectin deposition in inflammatory regions after EAE and MS (Haist et al., 2012; Sobel, 1998; Teesalu et al., 2001; van Horssen et al., 2006; van Horssen, Bo, Vos, Virtanen, & de Vries, 2005). To investigate the relationship between fibronectin deposition and fibroblast infiltration, we immunostained thoracic spinal cord tissue sections from Col1α1^GFP^ mice after EAE with an antibody against fibronectin. At the onset phase of EAE, there were small areas of fibronectin deposition that were closely associated with inflammatory regions but not with fibroblasts (Fig. 3F-J, U, W). However, at the acute and chronic phases, 70-85% of fibrotic areas were associated with fibronectin deposition (Fig. 3U) and 60-70% of the fibronectin positive area was associated with fibroblasts (Fig. 3W). Taken together, our data demonstrates that the fibrotic scar is closely associated with regions of inflammation and ECM deposition after EAE.

### Fibroblasts do not support OPC survival or promote OPC differentiation into oligodendrocytes

OPCs are present in fibrotic regions after EAE (Supplementary Fig. 3; (Tripathi, Rivers, Young, Jamen, & Richardson, 2010)) and MS (Chang, Nishiyama, Peterson, Prineas, & Trapp, 2000), but the effects of fibrosis on the regenerative capacity of OPCs is unknown. To investigate the effects of fibroblasts on OPC survival and differentiation, we performed a series of *in vitro* co-culture assays. OPCs will undergo apoptosis *in vitro* without certain growth factors (Barres et al., 1992) and have been shown to differentiate into oligodendrocytes upon growth factor withdrawal (Cheng et al., 2007; Tang, Tokumoto, & Raff, 2000). Cytopainter-labeled OPCs were plated on either fibroblast, astrocyte, or combined fibroblast and astrocyte monolayers and their response to growth factor withdrawal (media lacking PDGFA and bFGF) was analyzed. Astrocyte monolayers were used as a positive cellular control since astrocytes have previously been shown to support OPC survival in response to growth factor withdrawal (Pang et al., 2013). Mixed fibroblast and astrocyte monolayers were used to test for combined or compensatory effects. A PDL only condition was used as a non-biological control that would not provide trophic or contact-mediated cues for survival or differentiation. After two days *in vitro* (one day of growth factor withdrawal), fewer cytopainted OPCs were observed on PDL and fibroblast monolayers compared to astrocyte monolayers or mixed fibroblast+astrocyte monolayers, (Fig. 4E-H, M). These results suggest that OPC survival or proliferation was enhanced by astrocytes but not by fibroblasts which is consistent with previous reports that astrocytes provide supportive growth factors such as PDGF, LIF, and CNTF (Barres, Schmid, Sendnter, & Raff, 1993; Gard, Burrell, Pfeiffer, Rudge, & Williams, 1995; Raff, Lillien, Richardson, Burne, & Noble, 1988).

To test whether the decreased number of OPCs observed on fibroblast monolayers was due to decreased proliferation and/or increased cell death, we assessed incorporation of EdU or a dead cell stain. The percent of EdU^+^ proliferating OPCs was not different between substrates (Fig. 4A-L, N), but the percent of OPCs that were dead or dying was significantly higher for fibroblast monolayers compared to astrocyte monolayers and mixed fibroblast+astrocyte monolayers (Fig. 5). Cell death was similar between fibroblast monolayers and PDL, suggesting that fibroblasts do not provide an environment favorable for OPC survival after growth factor withdrawal.

To test whether fibroblasts affect the differentiation of OPCs into oligodendrocytes, co-cultures were fixed and stained for MBP after four days *in vitro* (three days of growth factor withdrawal). The percent of cytopainted OPCs that differentiated into MBP^+^ oligodendrocytes was significantly less on fibroblast monolayers and PDL compared to astrocyte monolayers (Fig. 6). Similar to cell death, the presence of supportive astrocytes in the mixed fibroblast+astrocyte condition is sufficient to rescue this effect although the mechanism remains unknown. Taken together, our data suggest that fibroblasts do not provide an environment conducive for OPC survival and differentiation.

### Fibroblast ECM and conditioned media inhibit the differentiation of OPCs into oligodendrocytes

The presence of fibroblasts in demyelinated lesions after EAE suggests they may be producing ECM molecules, cytokines, and/or growth factors that inhibit the differentiation of OPCs into remyelinating oligodendrocytes. To test whether fibroblast ECM inhibits the differentiation of OPCs, we cultured OPCs on fibroblast ECM and stimulated their differentiation with triiodothyronine (T_3_) and CNTF (Fig. 7A). To generate fibroblast ECM, we cultured primary meningeal fibroblasts on PDL and then performed decellularization using a commercially available non-enzymatic decellularization buffer (see methods). The presence of ECM was verified by immunostaining for fibronectin as a surrogate marker of ECM deposition, and proper decellularization was verified by the absence of DAPI stained nuclei (Fig. 7B-C). Approximately 33% of OPCs differentiated into MBP^+^ oligodendrocytes on PDL while only 22% differentiated on fibroblast ECM (Fig. 7D-F), suggesting that fibroblast ECM is inhibitory to oligodendrogenesis even in the presence of differentiation factors.

To test the paracrine effects of fibroblasts on OPC differentiation, we cultured OPCs in conditioned media supplemented with T_3_ and CNTF to stimulate their differentiation into oligodendrocytes (Fig. 8A). In non-conditioned media containing T_3_ and CNTF, approximately 30% of OPCs differentiate into MBP^+^ oligodendrocytes after three days (Fig. 8B, E). A similar and slightly increased percentage of OPCs differentiated in astrocyte conditioned media (35%) as expected based on our results in Figure 6, while significantly fewer OPCs differentiated into oligodendrocytes in fibroblast conditioned media (16%) (Fig. 8C-E). Taken together, our data suggest that fibroblasts actively inhibit the differentiation of OPCs into mature oligodendrocytes through the production of inhibitory soluble factors and/or ECM.

## Discussion

The extracellular environment of MS lesions plays a complex role in the regulation of inflammation and the regenerative capacity of oligodendrocyte progenitor cells. Targeting the sclerotic lesion environment for therapeutic purposes requires a better understanding of the cell populations that contribute to scar formation in MS. While astrogliosis in response to leukocyte infiltration has been the primary focus of research on the extracellular environment in most neurological disorders, our study identifies fibroblasts as another cell type that is abundant in demyelinating lesions after EAE. Fibroblasts are the principal cellular mediators of ECM deposition during wound healing in most tissues, but the extent to which fibroblasts contribute to the pathology of neuroinflammatory diseases such as MS has not previously been investigated. We show here that fibroblast activation coincides with the onset of EAE in the spinal cord and the optic nerve, and their peak of infiltration into spinal cord white matter occurs at the peak of motor deficits. During the chronic phase of EAE, much of the fibrotic area has resolved corresponding to decreases in CD11b^+^ myeloid cell accumulation and demyelinated area. Furthermore, we found that fibroblast conditioned media and fibroblast ECM inhibit the differentiation of OPCs into mature oligodendrocytes even without stimulating fibroblasts with pro-inflammatory cytokines, which highlights fibroblasts as potential mediators of an inhibitory environment in MS.

The appearance of inflammatory regions before fibroblast infiltration during the onset phase of EAE suggests that fibroblasts are proliferating/migrating into the CNS parenchyma in response to inflammatory leukocyte signaling. This hypothesis is supported by our previous study demonstrating that depletion of hematogenous macrophages reduces fibroblast infiltration after SCI (Zhu et al., 2015). Interestingly, while fibroblasts persist chronically after SCI (Soderblom et al., 2013), we observed a partial resolution of fibrotic regions chronically after EAE. This observation is consistent with decreased infiltration of myeloid cells after the acute phase (Supplementary Fig. 4), which suggests that the chronic persistence of fibroblasts after SCI may be due to continued infiltration of myeloid cells. These possibilities remain to be tested in the future. Notably, partial fibrotic resolution also corresponds to decreased demyelinated area and improved locomotor behavior suggesting a close association between fibrosis, demyelination, and paralysis, but the cause and effect relationships between these processes remain to be determined.

Due to the location of the fibrotic regions at the junction of blood vessels and meninges, it was difficult to distinguish the origin of fibroblasts in EAE. However, based on our previous study on SCI (Soderblom et al., 2013), it seems likely that a large subpopulation of the fibroblasts are of perivascular origin. Using genetic lineage tracing, we have previously shown that these perivascular fibroblasts are distinct from classical NG2-positive pericytes (Soderblom et al., 2013). Whereas pericytes are typically located around capillaries, perivascular fibroblasts are located preferentially along large diameter vessels, especially where they enter or leave the spinal cord parenchyma. Interestingly, this is the location of the Virchow-Robin space where leukocytes are able to enter from circulation. Accordingly, this region has been implicated as a trigger for inflammation during MS and EAE (Kawakami, Bartholomaus, Pesic, & Mues, 2012; Ransohoff & Engelhardt, 2012), and there is significant crosstalk between fibroblasts and leukocytes during inflammation (Van Linthout, Miteva, & Tschope, 2014) raising the possibility that fibroblasts in this perivascular/meningeal area may actively contribute to the inflammatory process.

The migration of fibroblasts to the injury site during wound healing is an important process for proper tissue regeneration, but excessive fibroblast migration/proliferation leads to detrimental tissue fibrosis. There have been conflicting reports as to whether increased ECM deposition is beneficial or detrimental to remyelination, likely owing to the complex dynamics of demyelinating models and the particular type of ECM protein under investigation (Baron, Shattil, & ffrench-Constant, 2002; De La Fuente et al., 2017; Maier et al., 2005; Stoffels et al., 2013; Zhao, Fancy, Franklin, & ffrench-Constant, 2009). The current leading hypothesis, however, is that aberrant ECM accumulation is inhibitory to the processes required for remyelination including the proliferation and migration of OPCs to the injury site as well as their differentiation into oligodendrocytes and eventual myelination of demyelinated axons. The data presented here supports this hypothesis and demonstrates that fibroblast ECM inhibits oligodendrogenesis. Additionally, we show that fibroblast conditioned media inhibits OPC differentiation, which may be due to the presence of inhibitory molecules, although the mechanisms underlying the effects of fibroblast ECM and conditioned media on oligodendrogenesis remain to be elucidated.

The Col1α1^GFP^ fibroblasts described here are one of several vessel-associated cell types that could be involved in the pathology of EAE and MS. For example, a recent study found that conditioned media from isolated pericytes can stimulate OPC differentiation, and pericytes (identified by PDGFRβ staining) proliferate in response to ethidium bromide-induced demyelination (De La Fuente et al., 2017). Another recent study of single-cell transcriptomics of brain vascular and vessel-associated cells identified “fibroblast-like” cells which specifically expressed *Col1a1* (Vanlandewijck et al., 2018). These cells are likely the same population labeled by the Col1α1^GFP^ transgenic mouse used here. Vanlandewijck et al. further demonstrated that the fibroblast-like cells were distinct from the pericyte population which highly expressed *Cspg4* (NG2), and the fibroblast, pericyte, and smooth muscle cell populations expressed *Pdgfrb*. We show here that Col1α1^GFP^ fibroblasts are positive for PDGFRβ (Supplementary Fig.1, (Soderblom et al., 2013)) which suggests that PDGFRβ may not be an appropriate marker to specifically identify pericytes. It is therefore possible that the documented increase in the number of pericytes in toxin-induced demyelinated lesions described by De La Fuente et al. may also include Col1α1 fibroblasts. Whether fibroblasts and pericytes have discrete functional roles in MS remains to be directly tested.

Overall, we have shown for the first time that fibroblasts contribute to EAE pathology by infiltrating the white matter parenchyma shortly after myeloid cells, and while fibrotic regions partially resolve over time, fibroblasts persist in areas of chronic demyelination. We further demonstrated that fibroblast ECM and fibroblast conditioned media inhibit the stimulated differentiation of OPCs into mature oligodendrocytes. Our data indicate that the fibrotic scar contributes to an environment that is inhibitory to remyelination in EAE lesions, and that future studies will need to investigate anti-fibrotic mechanisms as novel therapeutic targets in MS especially in combination with therapies that aim to promote OPC differentiation.

## Abbreviations

CNS: central nervous system
CNTF: ciliary neurotrophic factor
Col1α1: collagen 1 alpha 1
EAE: experimental autoimmune encephalomyelitis
ECM: extracellular matrix
EdU: 5-ethynyl-2’-deoxyuridine
GFP: green fluorescent protein
GFAP: glial fibrillary acidic protein
GLAST: glutamate aspartate transporter
LIF: leukemia inhibitory factor
MBP: myelin basic protein
MS: multiple sclerosis
OPC: oligodendrocyte progenitor cells
PDGFR: platelet derived growth factor receptor
SCI: spinal cord injury

## Acknowledgements

This study was funded by National MS Society PP-1510-06517, NINDS R01NS081040, R21NS082835, The Miami Project to Cure Paralysis, and the Buoniconti Fund. We thank Dr. David Brenner and Dr. Tatiana Kisseleva for the Col1α1^GFP^ transgenic mice. We thank Natasha Cammer, Eric Bray, Yadira Salgueiro and Shaffiat Karmally for administrative and technical assistance.

## References

Baron, W., Shattil, S. J., & ffrench-Constant, C. (2002). The oligodendrocyte precursor mitogen PDGF stimulates proliferation by activation of alpha(v)beta3 integrins. EMBO J, 21(8), 1957–1966. doi:10.1093/emboj/21.8.1957

Barres, B. A., Hart, I. K., Coles, H. S., Burne, J. F., Voyvodic, J. T., Richardson, W. D., & Raff, M. C. (1992). Cell death and control of cell survival in the oligodendrocyte lineage. Cell, 70(1), 31–46.

Barres, B. A., Schmid, R., Sendnter, M., & Raff, M. C. (1993). Multiple extracellular signals are required for long-term oligodendrocyte survival. Development, 118(1), 283–295.

Chang, A., Nishiyama, A., Peterson, J., Prineas, J., & Trapp, B. D. (2000). NG2-positive oligodendrocyte progenitor cells in adult human brain and multiple sclerosis lesions. J Neurosci, 20(17), 6404–6412.

Cheng, X., Wang, Y., He, Q., Qiu, M., Whittemore, S. R., & Cao, Q. (2007). Bone morphogenetic protein signaling and olig1/2 interact to regulate the differentiation and maturation of adult oligodendrocyte precursor cells. Stem Cells, 25(12), 3204–3214. doi:10.1634/stemcells.2007-0284

De La Fuente, A. G., Lange, S., Silva, M. E., Gonzalez, G. A., Tempfer, H., van Wijngaarden, P., … Rivera, F. J. (2017). Pericytes Stimulate Oligodendrocyte Progenitor Cell Differentiation during CNS Remyelination. Cell Rep, 20(8), 1755–1764. doi:10.1016/j.celrep.2017.08.007

Dias, D. O., & Goritz, C. (2018). Fibrotic scarring following lesions to the central nervous system. Matrix Biol. doi:10.1016/j.matbio.2018.02.009

Fernandez-Klett, F., & Priller, J. (2014). The fibrotic scar in neurological disorders. Brain Pathol, 24(4), 404–413. doi:10.1111/bpa.12162

Gard, A. L., Burrell, M. R., Pfeiffer, S. E., Rudge, J. S., & Williams, W. C., 2nd. (1995). Astroglial control of oligodendrocyte survival mediated by PDGF and leukemia inhibitory factor-like protein. Development, 121(7), 2187–2197.

Haist, V., Ulrich, R., Kalkuhl, A., Deschl, U., & Baumgartner, W. (2012). Distinct spatio-temporal extracellular matrix accumulation within demyelinated spinal cord lesions in Theiler’s murine encephalomyelitis. Brain Pathol, 22(2), 188–204. doi:10.1111/j.1750-3639.2011.00518.x

Hibbits, N., Yoshino, J., Le, T. Q., & Armstrong, R. C. (2012). Astrogliosis during acute and chronic cuprizone demyelination and implications for remyelination. ASN Neuro, 4(6), 393–408. doi:10.1042/AN20120062

Kawakami, N., Bartholomaus, I., Pesic, M., & Mues, M. (2012). An autoimmunity odyssey: how autoreactive T cells infiltrate into the CNS. Immunol Rev, 248(1), 140–155. doi:10.1111/j.1600-065X.2012.01133.x

Lau, L. W., Cua, R., Keough, M. B., Haylock-Jacobs, S., & Yong, V. W. (2013). Pathophysiology of the brain extracellular matrix: a new target for remyelination. Nat Rev Neurosci, 14(10), 722–729. doi:10.1038/nrn3550

Maier, O., van der Heide, T., van Dam, A. M., Baron, W., de Vries, H., & Hoekstra, D. (2005). Alteration of the extracellular matrix interferes with raft association of neurofascin in oligodendrocytes. Potential significance for multiple sclerosis? Mol Cell Neurosci, 28(2), 390–401. doi:10.1016/j.mcn.2004.09.012

Mayo, L., Trauger, S. A., Blain, M., Nadeau, M., Patel, B., Alvarez, J. I., … Quintana, F. J. (2014). Regulation of astrocyte activation by glycolipids drives chronic CNS inflammation. Nat Med, 20(10), 1147–1156. doi:10.1038/nm.3681

Mohan, H., Krumbholz, M., Sharma, R., Eisele, S., Junker, A., Sixt, M., … Meinl, E. (2010). Extracellular matrix in multiple sclerosis lesions: Fibrillar collagens, biglycan and decorin are upregulated and associated with infiltrating immune cells. Brain Pathol, 20(5), 966–975. doi:10.1111/j.1750-3639.2010.00399.x

Pang, Y., Fan, L. W., Tien, L. T., Dai, X., Zheng, B., Cai, Z., … Bhatt, A. (2013). Differential roles of astrocyte and microglia in supporting oligodendrocyte development and myelination in vitro. Brain Behav, 3(5), 503–514. doi:10.1002/brb3.152

Raff, M. C., Lillien, L. E., Richardson, W. D., Burne, J. F., & Noble, M. D. (1988). Platelet-derived growth factor from astrocytes drives the clock that times oligodendrocyte development in culture. Nature, 333(6173), 562–565. doi:10.1038/333562a0

Ransohoff, R. M., & Engelhardt, B. (2012). The anatomical and cellular basis of immune surveillance in the central nervous system. Nat Rev Immunol, 12(9), 623–635. doi:10.1038/nri3265

Sobel, R. A. (1998). The extracellular matrix in multiple sclerosis lesions. J Neuropathol Exp Neurol, 57(3), 205–217.

Sobel, R. A., & Ahmed, A. S. (2001). White matter extracellular matrix chondroitin sulfate/dermatan sulfate proteoglycans in multiple sclerosis. J Neuropathol Exp Neurol, 60(12), 1198–1207.

Soderblom, C., Luo, X., Blumenthal, E., Bray, E., Lyapichev, K., Ramos, J., … Lee, J. K. (2013). Perivascular fibroblasts form the fibrotic scar after contusive spinal cord injury. J Neurosci, 33(34), 13882–13887. doi:10.1523/jneurosci.2524-13.2013

Stoffels, J. M., de Jonge, J. C., Stancic, M., Nomden, A., van Strien, M. E., Ma, D., … Baron, W. (2013). Fibronectin aggregation in multiple sclerosis lesions impairs remyelination. Brain, 136(Pt 1), 116–131. doi:10.1093/brain/aws313

Stoffels, J. M., Hoekstra, D., Franklin, R. J., Baron, W., & Zhao, C. (2015). The EIIIA domain from astrocyte-derived fibronectin mediates proliferation of oligodendrocyte progenitor cells following CNS demyelination. Glia, 63(2), 242–256. doi:10.1002/glia.22748

Tang, D. G., Tokumoto, Y. M., & Raff, M. C. (2000). Long-term culture of purified postnatal oligodendrocyte precursor cells. Evidence for an intrinsic maturation program that plays out over months. J Cell Biol, 148(5), 971–984.

Teesalu, T., Hinkkanen, A. E., & Vaheri, A. (2001). Coordinated induction of extracellular proteolysis systems during experimental autoimmune encephalomyelitis in mice. Am J Pathol, 159(6), 2227–2237. doi:10.1016/s0002-9440(10)63073-8

Tripathi, R. B., Rivers, L. E., Young, K. M., Jamen, F., & Richardson, W. D. (2010). NG2 glia generate new oligodendrocytes but few astrocytes in a murine experimental autoimmune encephalomyelitis model of demyelinating disease. J Neurosci, 30(48), 16383–16390. doi:10.1523/jneurosci.3411-10.2010

van Horssen, J., Bo, L., Dijkstra, C. D., & de Vries, H. E. (2006). Extensive extracellular matrix depositions in active multiple sclerosis lesions. Neurobiol Dis, 24(3), 484–491. doi:10.1016/j.nbd.2006.08.005

van Horssen, J., Bo, L., Vos, C. M., Virtanen, I., & de Vries, H. E. (2005). Basement membrane proteins in multiple sclerosis-associated inflammatory cuffs: potential role in influx and transport of leukocytes. J Neuropathol Exp Neurol, 64(8), 722–729.

Van Linthout, S., Miteva, K., & Tschope, C. (2014). Crosstalk between fibroblasts and inflammatory cells. Cardiovasc Res, 102(2), 258–269. doi:10.1093/cvr/cvu062

Vanlandewijck, M., He, L., Mae, M. A., Andrae, J., Ando, K., Del Gaudio, F., … Betsholtz, C. (2018). A molecular atlas of cell types and zonation in the brain vasculature. Nature, 554(7693), 475–480. doi:10.1038/nature25739

Yata, Y., Scanga, A., Gillan, A., Yang, L., Reif, S., Breindl, M., … Rippe, R. A. (2003). DNase I-hypersensitive sites enhance alpha1(I) collagen gene expression in hepatic stellate cells. Hepatology, 37(2), 267–276. doi:10.1053/jhep.2003.50067

Zhao, C., Fancy, S. P., Franklin, R. J., & ffrench-Constant, C. (2009). Up-regulation of oligodendrocyte precursor cell alphaV integrin and its extracellular ligands during central nervous system remyelination. J Neurosci Res, 87(15), 3447–3455. doi:10.1002/jnr.22231

Zhu, Y., Soderblom, C., Krishnan, V., Ashbaugh, J., Bethea, J. R., & Lee, J. K. (2015). Hematogenous macrophage depletion reduces the fibrotic scar and increases axonal growth after spinal cord injury. Neurobiol Dis, 74, 114–125. doi:10.1016/j.nbd.2014.10.024

