## Supplemental Figures for "Fibrotic scar after experimental autoimmune encephalomyelitis inhibits oligodendrocyte differentiation"

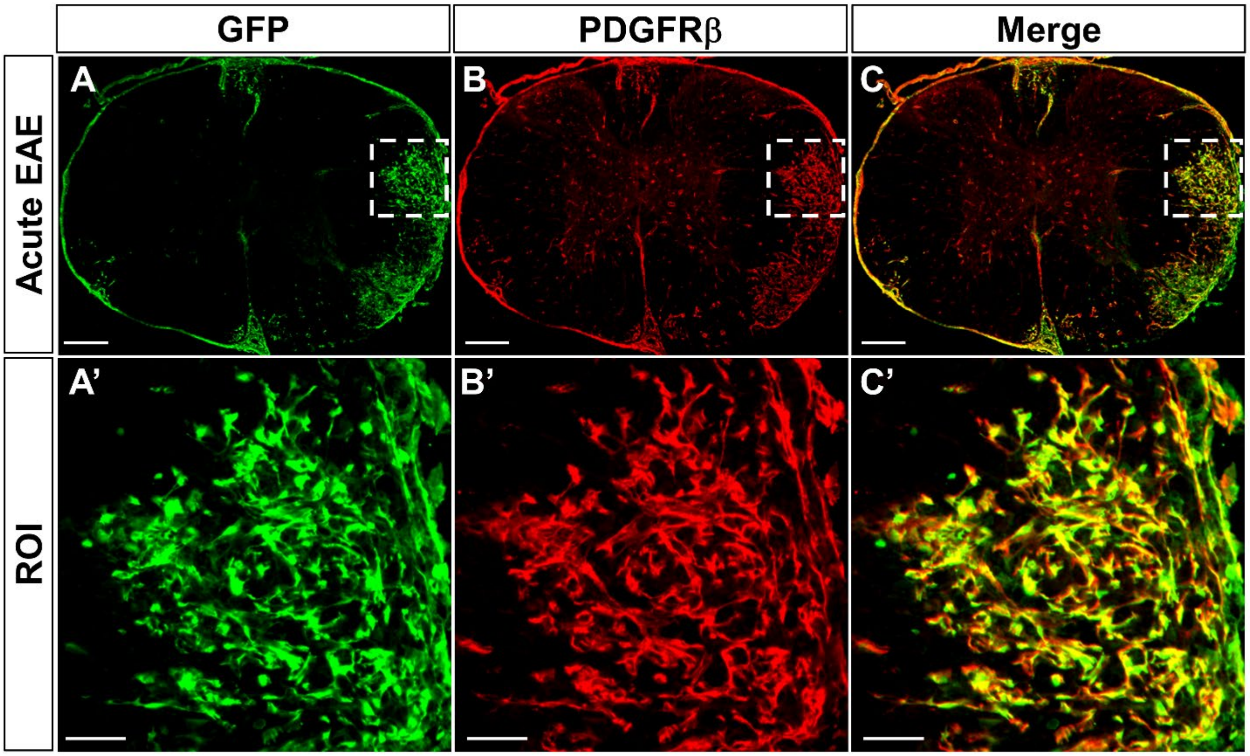


**Supplementary Figure 1.** Col1α1$^{GFP}$ fibroblasts in EAE lesions are PDGFRβ positive. Infiltrating Col1a1$^{GFP}$ fibroblasts colocalize with PDGFRβ immunostaining in the thoracic spinal cord during the acute phase of EAE. We had previously identified PDGFRβ as a marker of perivascular fibroblasts after spinal cord injury (Soderblom et al., 2013). Scale bars = 200μm (A-C) and 50μm (A'-B').

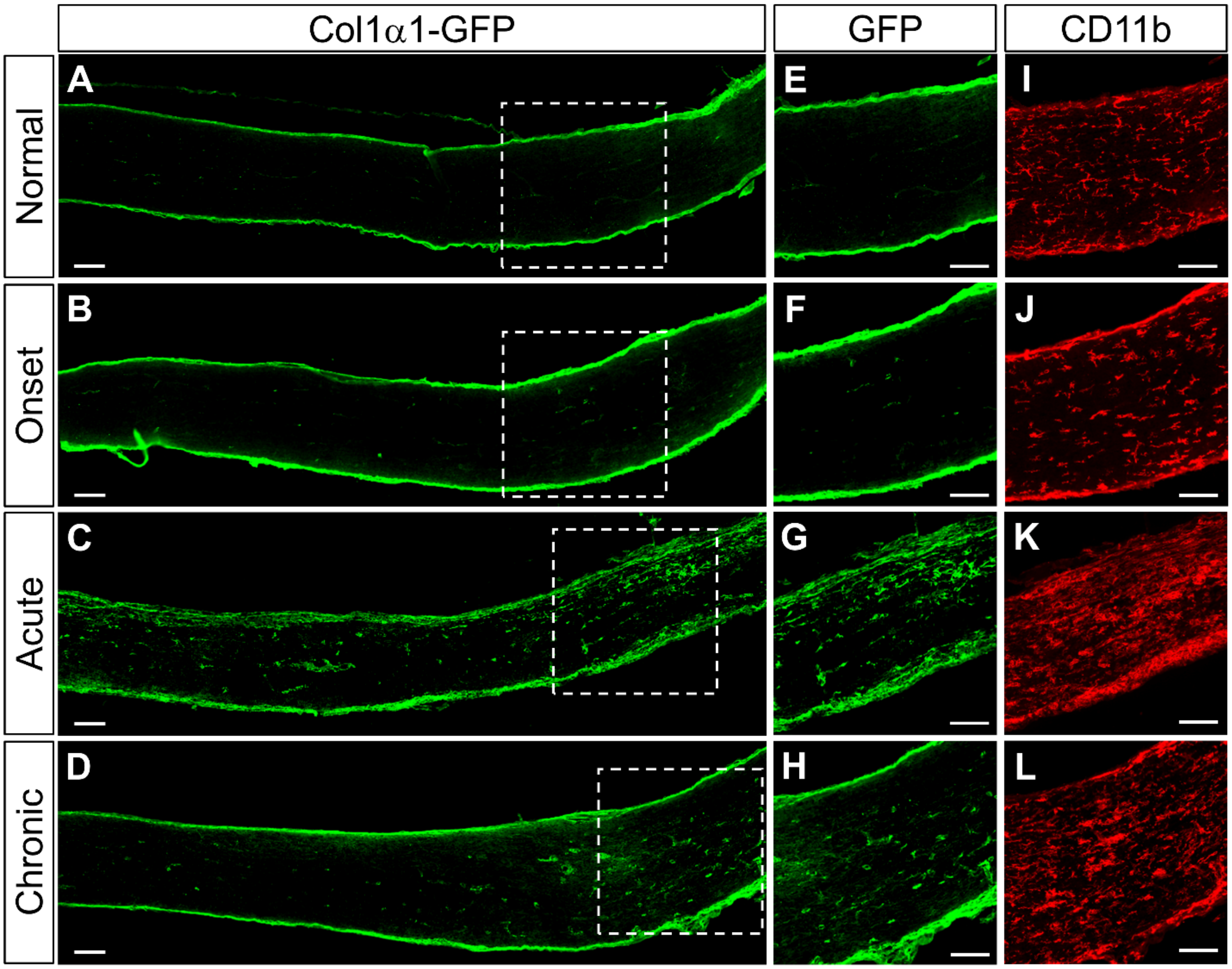


**Supplementary Figure 2**. Fibroblast activation also occurs in the optic nerve after EAE. Similar to the spinal cord, the optic nerve from a normal Col1α1$^{GFP}$ mouse shows fibroblasts in the meninges and around blood vessels (A, E). Fibroblasts appear normal at disease onset (B, F) and become reactive during acute disease (C, G) corresponding to an increase in CD11b+ macrophages/microglia (I-K). Fibroblast activation resolves chronically but not to normal levels (D, H). Scale bars = 100μm.

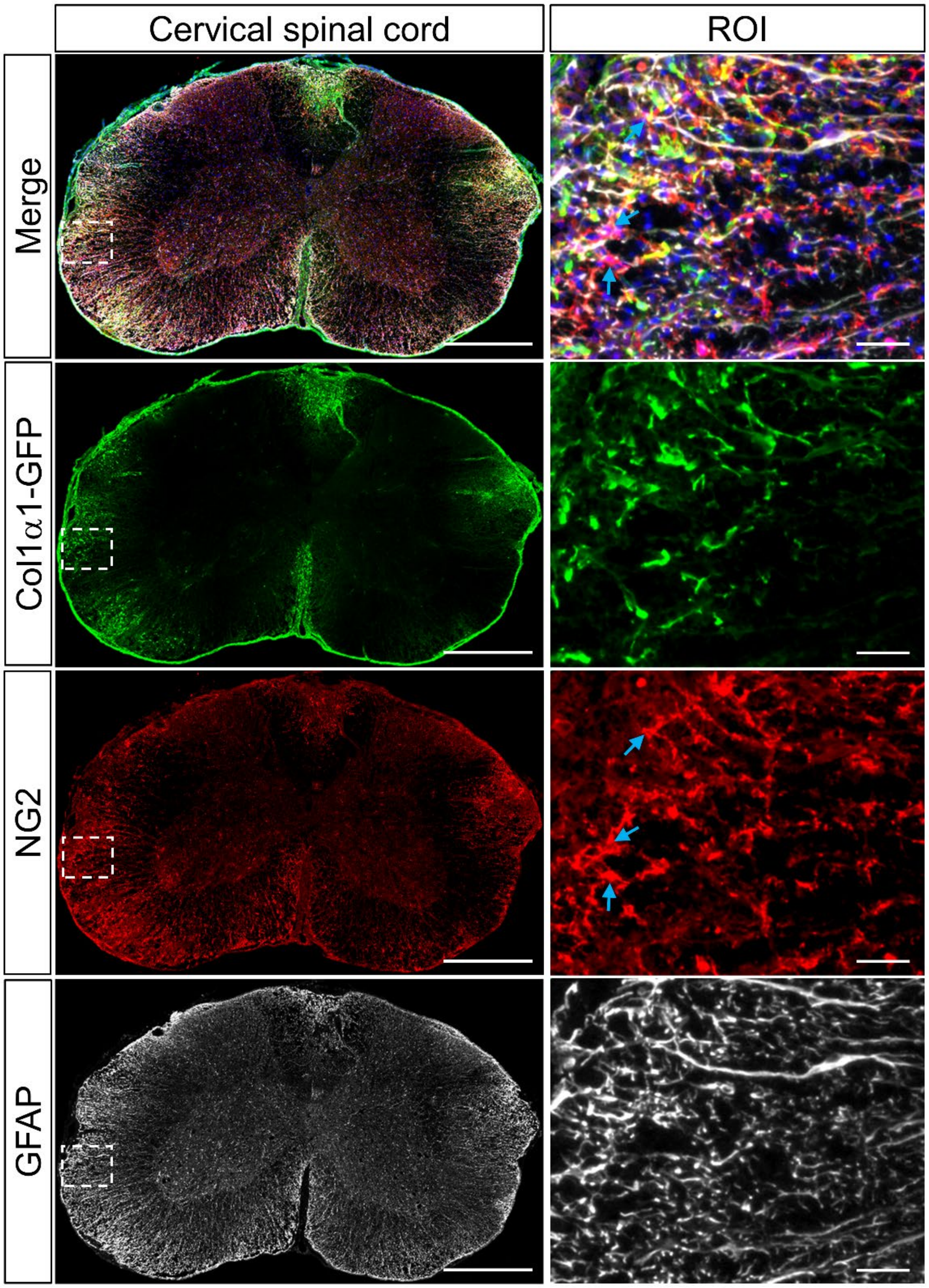


**Supplementary Figure 3.** OPCs and astrocytes are present in fibrotic regions after EAE. $GFAP^+$ astrocytes and $NG2^+$ OPCs (arrows) are present in areas with $Col1\alpha1^{GFP}$ fibroblasts in EAE lesions. Region of interest (ROI) is denoted by boxed regions. Scale bars = 500μm and 50μm (ROI).

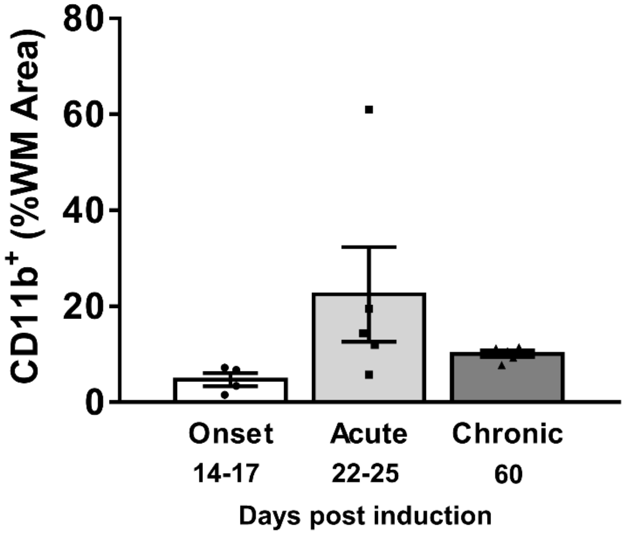


**Supplementary Figure 4**. Macrophage infiltration is reduced chronically after EAE. The percent of white matter that has a dense accumulation of $CD11b^+$ macrophages increases as the disease progresses from the onset to acute phase, and tends to decrease chronically
